# Dental and facial characteristics of osteogenesis imperfecta type V

**DOI:** 10.1101/413633

**Authors:** Jean-Marc Retrouvey, Doaa Taqi, Faleh Tamimi, Didem Dagdeviren, Francis H. Glorieux, Brendan Lee, Renna Hazboun, Deborah Krakow, V. Reid Sutton, Members of the BBD Consortium, Michael Bober, Paul Esposito, David R Eyre, Danielle Gomez, Gerald Harris, Tracy Hart, Mahim Jain, Jeffrey Krisher, Sandesh CS Nagamani, Eric S Orwoll, Cathleen L Raggio, Eric Rush, Peter Smith, Laura Tosi, Frank Rauch

## Abstract

Osteogenesis imperfecta (OI) type V is an ultrarare heritable bone disorder caused by the heterozygous c.-14C>T mutation in *IFITM5*. The dental and craniofacial phenotype has not been described in detail. In the present multicenter study (Brittle Bone Disease Consortium) 14 individuals with OI type V (age 3 to 50 years; 10 females, 4 males) underwent dental and craniofacial assessment. None of the individuals had dentinogenesis imperfecta. Six of the 9 study participants (66%) for whom panoramic radiographs were obtained had at least one missing tooth (range 1 to 9). Class II molar occlusion was present in 8 (57%) of the 14 study participants. The facial profile was retrusive and lower face height was decreased in 8 (57%) individuals. Cephalometry, performed in three study participants, revealed a severely retrusive maxilla and mandible, and poorly angulated incisors in a 14-year old girl, a protrusive maxilla and a retrusive mandible in a 14-year old boy. Cone beam computed tomograpy scans were obtained from two study participants and demonstrated intervertebral disc calcification at the C2-C3 level in one individual. Our study observed that OI type V is associated with missing permanent teeth, especially permanent premolar, but not with dentinogenesis imperfecta. The pattern of craniofacial abnormalities in OI type V thus differs from that in other severe OI types, such as OI type III and IV, and could be described as a bimaxillary retrusive malocclusion with reduced lower face height and multiple missing teeth.

## Introduction

Osteogenesis imperfecta (OI) is a heritable connective tissue disorder that is associated with bone fragility and often short stature (1). About 90% of individuals with a clinical diagnosis of OI have variants in *COL1A1* and *COL1A2*, the genes encoding collagen type I alpha chains (2). Clinically, such patients are classified into OI types I (least severe form), OI type II (perinatal lethal), OI type III (severe), OI type IV (moderate severity). There is presently no cure for OI, but bisphosphonate drugs are widely administered to increase bone density and decrease the number of fractures (1). Apart from fractures in long bones and vertebra, individuals with *COL1A1*- and *COL1A2*–related OI often have dental and craniofacial abnormalities, including dentinogenesis imperfecta (DI), tooth agenesis and malocclusion (3-5).

Among OI types not caused by *COL1A1* or *COL1A2*, OI type V is the most prevalent disorder (6). OI type V is caused by the recurrent heterozygous c.-14C>T variant in *IFITM5*, which encodes BRIL, a transmembrane protein with unknown function that is specifically expressed in osteoblasts (7-9). The variant creates a new translational start site and leads to the addition of 5 amino acids at the N-terminus of the BRIL protein. OI type V resembles OI type IV with regard to fracture incidence, long-bone deformities, vertebral compression fractures and scoliosis (10, 11), but also has distinguishing features such as hyperplastic callus formation and calcification of the interosseous forearm membrane (10, 12). How the addition of 5 amino acids to BRIL protein leads to bone fragility is unknown at present. Mouse models harboring the OI type V variant die at birth, which complicates mechanistic studies (13, 14).

Although OI type V seems to lead to similar bone material abnormalities and bone fragility as OI type IV caused by *COL1A1*/*COL1A2* mutations (10, 15), little is known about the involvement of the craniofacial skeleton and teeth in OI type V. Previous reports on OI type V stated that affected individuals did not have clinical signs of DI (10, 16, 17), but missing teeth, short roots and ectopic eruption of the molars have been reported (17). A prospective, detailed characterization of the craniofacial and dental phenotype in a cohort of OI type V has not been performed.

In the present report we therefore evaluated the dental and craniofacial characteristics in a cohort of individuals with OI type V who were identified through the Brittle Bone Disease Consortium, a multicenter Rare Disease Clinical Research Network.

## Materials and Methods

### Study Participants

Individuals with a diagnosis of OI type V were recruited through the Brittle Bone Disease Consortium(https://www.rarediseasesnetwork.org/cms/BBD) that comprises several specialized centers from across North America (Houston, Montreal, Chicago, Baltimore, Portland, Washington DC, New York, Omaha, Los Angeles). One of the projects conducted by the consortium is a natural history study to assess the clinical features of OI. Patients with a diagnosis of OI of any type and any age are eligible to participate. Dental evaluation is offered to participants who are three years of age or older.

The present study analyzes baseline dental data from the 14 study participants (10 females, 4 males; age range 3 to 50 years) who had a diagnosis of OI type V. Two individuals in addition agreed to participate in an ancillary study involving cone beam computed tomography (CBCT). For 11 individuals, the diagnosis of OI type V was confirmed by genetic testing (heterozygosity for the c.-14C>T variant in *IFITM5*); in 3 participants the diagnosis was based on clinical findings alone. Three cephalometric radiographs were also studied. The study was approved at all participating study centers, and all study participants or their legal guardians provided informed consent.

### Craniofacial and dental evaluations

The dental evaluation comprised clinical examination and extra- and intraoral photographs for all study participants, panoramic radiographs for those participants aged six years and older who consented to the test. The study dentist at each site performed the intraoral clinical examination and was responsible for obtaining a standard set of intraoral and extraoral photographs as well as panoramic radiographs. Examiners had been calibrated by passing the simplified International Caries Detection and Assessment System (ICDAS) examination (18). The photographs and radiographs were uploaded to the study website and were independently assessed by two experienced central readers, an orthodontist and an oral radiologist. There were no inter-observer disagreements on the assessments presented here. CBCT scans were evaluated by an oral and maxillofacial radiologist.

The clinical dental examination included assessing the classification of occlusion, overbite, overjet, crowding, open bite, crossbite, arch shape and the presence of DI (said to be present if at least one tooth appearing opalescent or had gray, brown or yellow color). The presence of DI was also assessed by examining panoramic radiographs (radiographic criteria for DI were: partial or complete pulp obliteration, bulbous crown, cervical constrictions, roots abnormalities or short root, and taurodontism).

Lateral and frontal photographs were used to assess facial type (normocephalic, brachycephalic or dolichocephalic; facial symmetry; facial proportions), and profile (normal, concave or convex), ear position (19), and frontal bossing. The facial type was determined by using the facial index that was calculated as the ratio between the maximum width to the maximum length of the face on the frontal photograph: dolichocephalic (long and narrow face), <0.72; normocephalic, 0.72 to 0.82; brachycephalic (short and wide face), >0.82 (20). Facial symmetry was determined by comparing the right and left sides of the face as determined by a reference midline passing through the glabella (midpoint between the eyebrow) and the subnasale points (the junction between the nasal septum and the upper lip). Facial proportions were calculated as follows: total face height, distance between glabella and soft tissue pogonion (the most anteroinferior point of the chin); upper face height, distance between glabella and subnasale; lower face height, distance between subnasale and the soft tissue pogonion. A lower face height less than 47% of total face height was classified as reduced; a lower face height more than 63% of total face height was classified as increased (21). Profile assessment was performed by measuring the angle between glabella, subnasale, and soft tissue pogonion in a full profile picture. An angle between 158 and 180 degrees was considered normal, regardless of sex (22). An angle <158 degrees indicates a convex profile while an angle >180 degree represents a flat to concave profile.

Lateral cephalometric radiographs were analyzed using the Ceph Tracing routine of the Dolphin Imaging software package (version 11.8). Results were compared to normative cephalometric data as published (23). CBCT scans were acquired with a 3D Accuitomo 170 (Morita Inc, Kyoto, Japan) CBCT machine in a 170 mm x 120 mm field-of-view and a 250 μm voxel size. The exposure settings for CBCT included a tube voltage of 90 kV and a tube current of 4.5 mA for 17.5 seconds. Image analysis was performed using Anatomage InVivo 5 version 5.4 (Invivo Dental; Anatomage, San Jose, CA) software.

In order to evaluate the projection of the lip and the convexity of the lower face, two measurements were used. The H angle measures the prominence of the lips in relation to the facial line (Nasion soft-pogonion soft: Normal mean value: 6 degrees, standard deviation 3 degrees) (24). The Z angle is used to measure facial balance and measures the prominence of the lips or the relative retrusiveness of the chin. The Z angle is formed by the intersection of the lines Na (soft tissue)-Pogonion (soft tissue) and Pogonion(soft tissue) - most prominent lip point. The normal mean for the Z angle is 10 degrees with a standard deviation of 4 degrees (25).

Taking into account frequent reports of hypodontia in patients with OI, to evaluate the caries prevalence in this population a modification of the DFT index was utilized. Wherein, caries scores were adjusted for the missing teeth, by dividing the total sum of decayed and missing teeth by the total number of teeth present in an individual. Therefore, this adjusted DFT is a continuous variable ranging from 0 to 1, with 0 being the minimum caries experience and 1 being the highest (26).

## Results

Fourteen study participants with OI type V underwent dental evaluation (Table 1). Panoramic radiographs were obtained in 9 study participants. None of the individuals had clinical or radiological signs of DI. Tooth discoloration, pulp obliteration, bulbous crowns, short roots or taurodontism were not observed in this cohort. Six of the nine study participants (55%) for whom panoramic radiographs were obtained had at least one missing tooth. The number of missing teeth ranged from 1 to 9. Three of these 9 individuals had retained deciduous teeth past the normal range of exfoliation; eight study participants had impacted teeth. Ten study participants had an adjusted DFT score of 0, in four subjects the adjusted DFT score ranged from 0.21 to 0.67. Examples of the clinical and radiological features are shown in Figures 1 and 2.

**Table 1.**
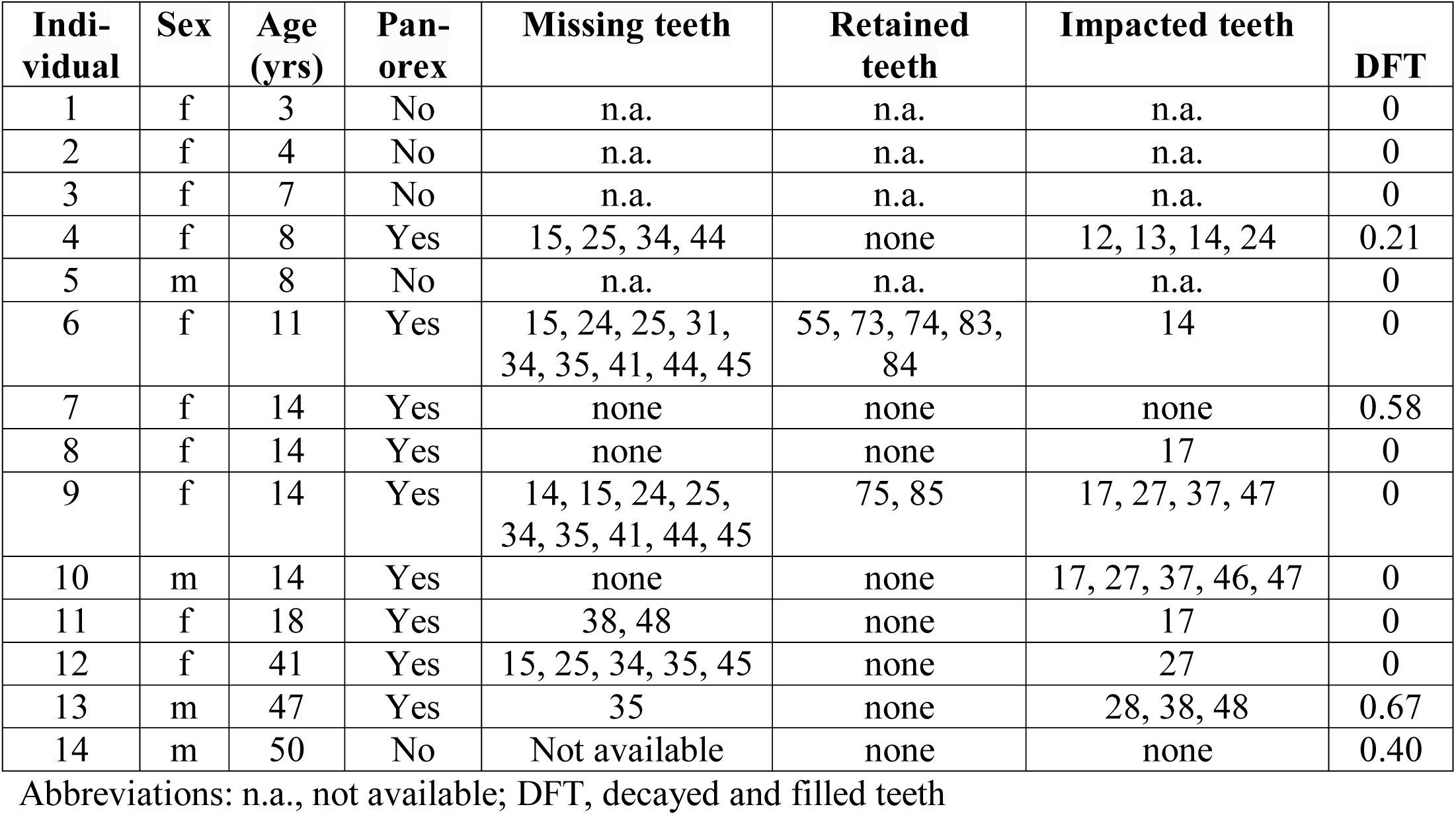
**General clinical information and dental evaluation.**

| **Indi-**<br>**vidual** | **Sex** | **Age**<br>**(yrs)** | **Pan-**<br>**orex** | **Missing teeth** | **Retained**<br>**teeth** | **Impacted teeth** | **DFT** |
| --- | --- | --- | --- | --- | --- | --- | --- |
| 1 | f | 3 | No | n.a. | n.a. | n.a. | 0 |
| 2 | f | 4 | No | n.a. | n.a. | n.a. | 0 |
| 3 | f | 7 | No | n.a. | n.a. | n.a. | 0 |
| 4 | f | 8 | Yes | 15, 25, 34, 44 | none | 12, 13, 14, 24 | 0.21 |
| 5 | m | 8 | No | n.a. | n.a. | n.a. | 0 |
| 6 | f | 11 | Yes | 15, 24, 25, 31,<br>34, 35, 41, 44, 45 | 55, 73, 74, 83,<br>84 | 14 | 0 |
| 7 | f | 14 | Yes | none | none | none | 0.58 |
| 8 | f | 14 | Yes | none | none | 17 | 0 |
| 9 | f | 14 | Yes | 14, 15, 24, 25,<br>34, 35, 41, 44, 45 | 75, 85 | 17, 27, 37, 47 | 0 |
| 10 | m | 14 | Yes | none | none | 17, 27, 37, 46, 47 | 0 |
| 11 | f | 18 | Yes | 38, 48 | none | 17 | 0 |
| 12 | f | 41 | Yes | 15, 25, 34, 35, 45 | none | 27 | 0 |
| 13 | m | 47 | Yes | 35 | none | 28, 38, 48 | 0.67 |
| 14 | m | 50 | No | Not available | none | none | 0.40 |

Abbreviations: n.a., not available; DFT, decayed and filled teeth

**Figure 1.**
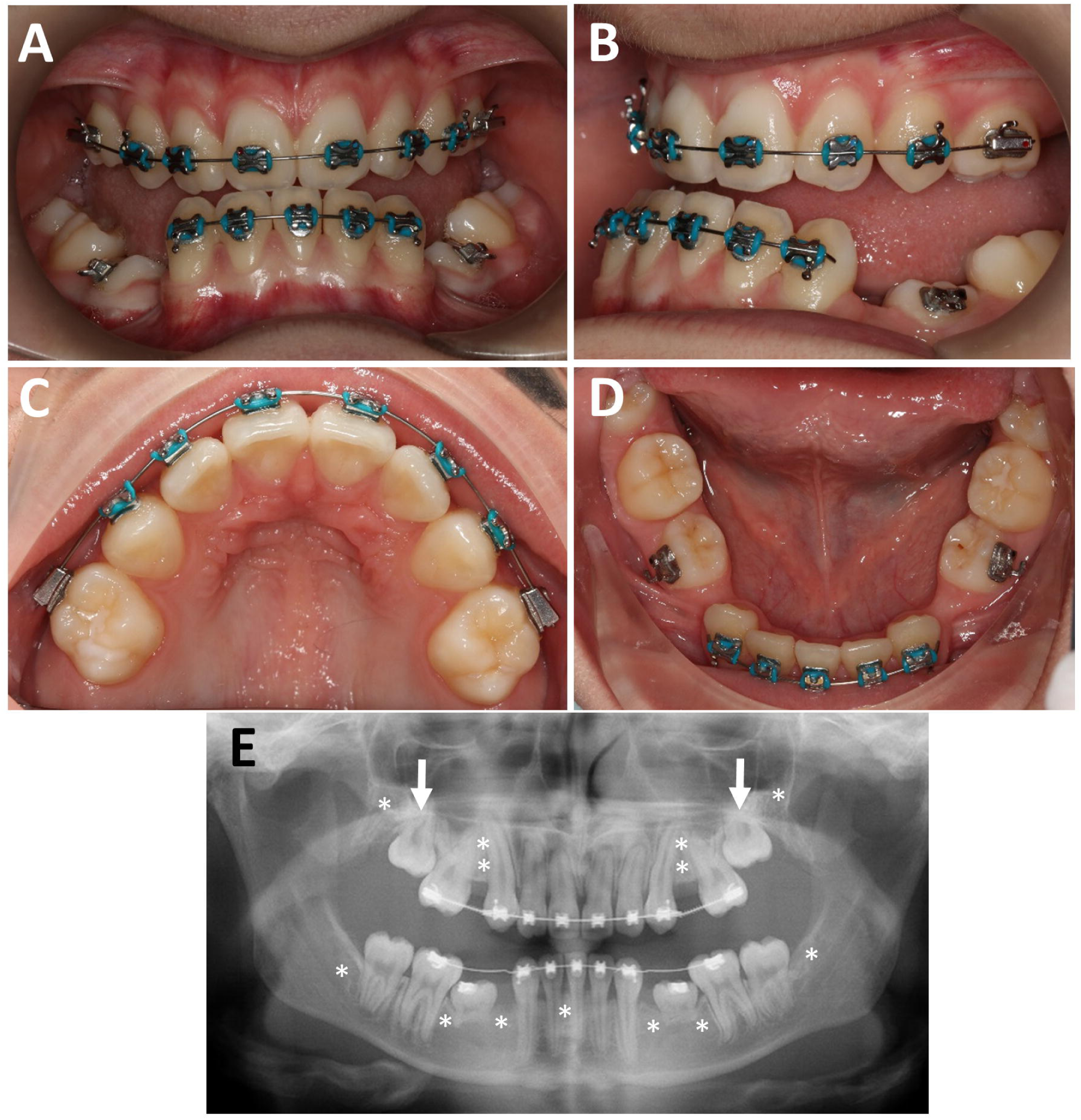
OI type V, Individual 9 (girl, 14 years). **A, B.** Intraoral photographs, showing posterior open bite, and Class III malocclusion. **C, D.** Occlusal view, showing missing premolars and retained lower deciduous molars. **E.** Panoramic radiograph, showing the deciduous molars with resorbed roots, unerupted 2^nd^ molar (arrows), a missing lower incisor and the position of the 8 missing premolars (asterisks). Third molars are also congenitally missing (asterisks).

**Figure 2.**
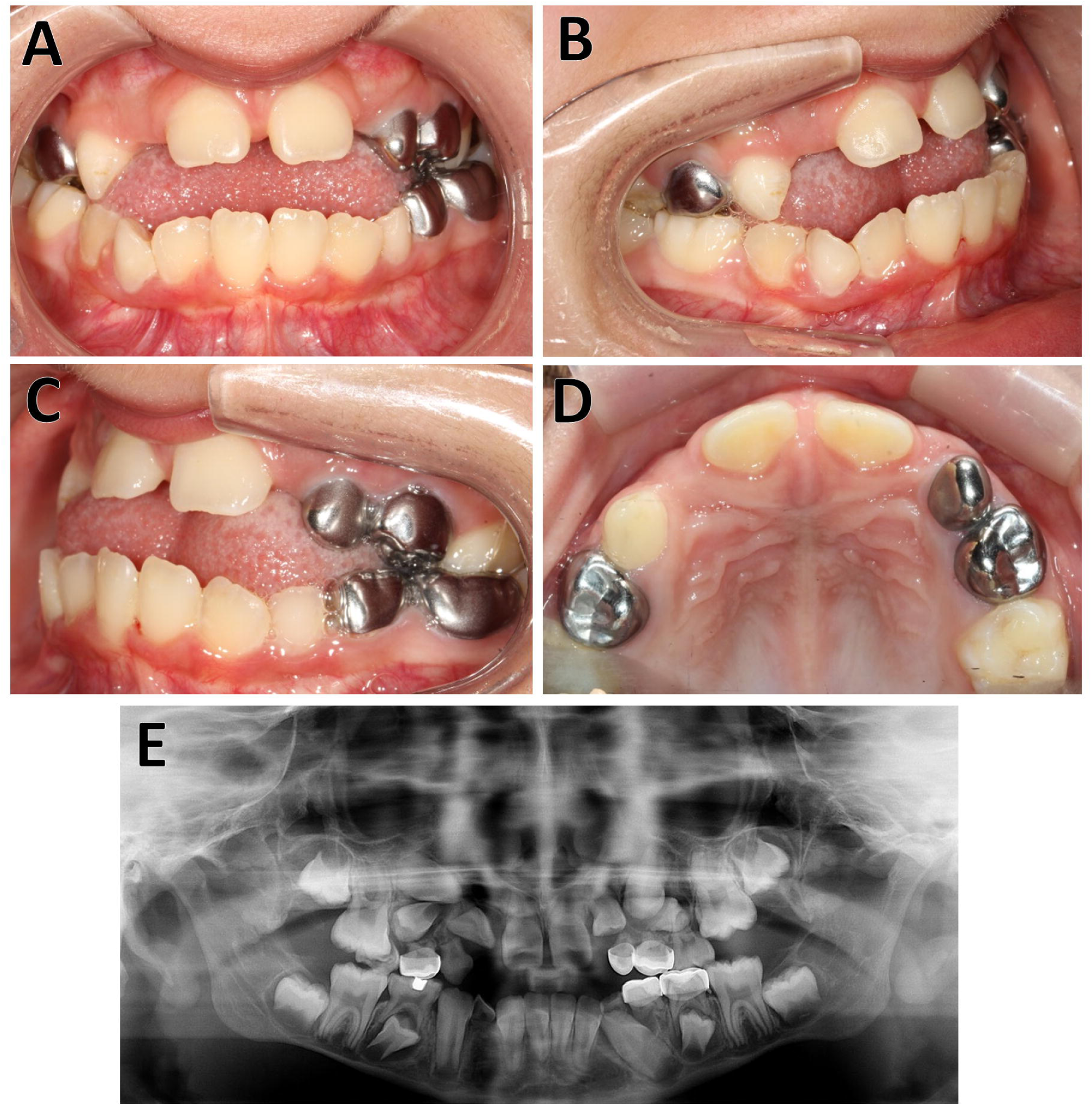
Individual 4 (girl, 8 years). **A, B, C** Intra-oral photographs, showing the anterior open bite with Class II malocclusion due to migration of upper permanent molars. **D.** Occlusal plane view. The radiograph shows the agenesis of the premolars (15, 25, 44, 34) as well as multiple impactions (12, 13,14, 24).

All study participants had normal ear position and no frontal bossing (Table 3). The facial profile was retrusive in 8 of the 14 study participants (57%). Assessment of facial proportions showed decreased lower face height in 8 individuals (57%).

**Table 2.** **Occlusion data.**

| Individual | Molar occl right | Molar occl left | Canine occl right | Canine occl left | Severity of mal-occlusion | Over-jet (mm) | Over-bite (%) | Anterior cross-bite | Posterior cross-bite | Anterior open bite | Lateral open bite | Crow-ding |
| --- | --- | --- | --- | --- | --- | --- | --- | --- | --- | --- | --- | --- |
| 1 | I | II | II | II | Mild | 2 | 15 | y | y | n | y | mild |
| 2 | II | I | II | I | severe | 0 | Openbite | y | y | y | y | n |
| 3 | I | I | I | I | mild | 1 | 0 | n | n | n | n | n |
| 4 | II | II | n.a. | n.a. | severe | -2 | Openbite | y | y | y | n | mild |
| 5 | II | II | n.a. | I | mild | 0 | Openbite | y | n | y | n | mild |
| 6 | II | II | n.a. | n.a. | severe | 4 | 100 | n | y | n | n | n |
| 7 | I | I | I | I | mild | 2 | 10 | n | n | n | n | n |
| 8 | II | II | II | II | moderate | 0 | Openbite | y | y | y | n | n |
| 9 | II | II | III | III | severe | -1 | 0 | y | y | y | y | n |
| 10 | II | II | II | II | severe | 6 | 40 | n | n | n | n | mild |
| 11 | I | I | I | I | n.a.<br>(braces) | 2 | 25 | n | n | n | n | n |
| 12 | I | I | I | I | moderate | 2 | 75 | n | n | n | n | moderate |
| 13 | I | III | I | II | severe | 0 | Openbite | y | y | y | y | n |
| 14 | III | III | III | II | mild | 2 | 25 | n | y | n | y | mild |

Abbreviation: n, no; n.a., not available; occl, occlusion; y, yes

**Table 3.** Facial characteristics.

| **Indi-**<br>**vidual** | **Face**<br>**Type** | **Facial**<br>**Profile** | **H line** | **Z angle** | **Facial**<br>**Proportions** | **Facial**<br>**Proportions (%)** |
| --- | --- | --- | --- | --- | --- | --- |
| 1 | brachy | retrusive | 7.8 | 76.9 | normal | 46 |
| 2 | brachy | retrusive | Na | n.a | NA | na |
| 3 | normo | normal | 19.1 | 70.4 | normal | 54 |
| 4 | normo | retrusive | 10.8 | 71.3 | decreased LFH | 45 |
| 5 | normo | normal | 24.5 | 59.3 | Normal | 52 |
| 6 | brachy | retrusive | 24.7 | 56.8 | Normal | 53 |
| 7 | brachy | retrusive | 3.7 | 78.3 | decreased LFH | 46 |
| 8 | normo | normal | 20.8 | 76.4 | Normal | 50 |
| 9 | brachy | retrusive | 0.1 | 91.8 | decreased LFH | 47 |
| 10 | brachy | convex | 24.6 | 65.8 | decreased LFH | 43 |
| 11 | brachy | retrusive | 11.2 | 76.8 | normal | 50 |
| 12 | normo | retrusive | 1.4 | 86.6 | decreased LFH | 43 |
| 13 | normo | normal | 6.1 | 88.6 | Normal | 47 |
| 14 | normo | normal | 9.3 | 99.9 | normal | 52 |

Abbreviations: LFH: lower face height; Normo: Normocephalic; Brachy: Brachycephalic

Cephalometry was performed in three study participants (Figure 3). Individual 9, a 14-year old girl, presented a severely retrusive maxilla and mandible, multiple missing teeth, an underdeveloped lower face and poorly angulated upper and lower incisors. In contrast, individual 10, a 14-year old boy, had a protrusive maxilla and a retrusive mandible. Individual 13 presented with almost normal cephalometric results.

**Figure 3.**
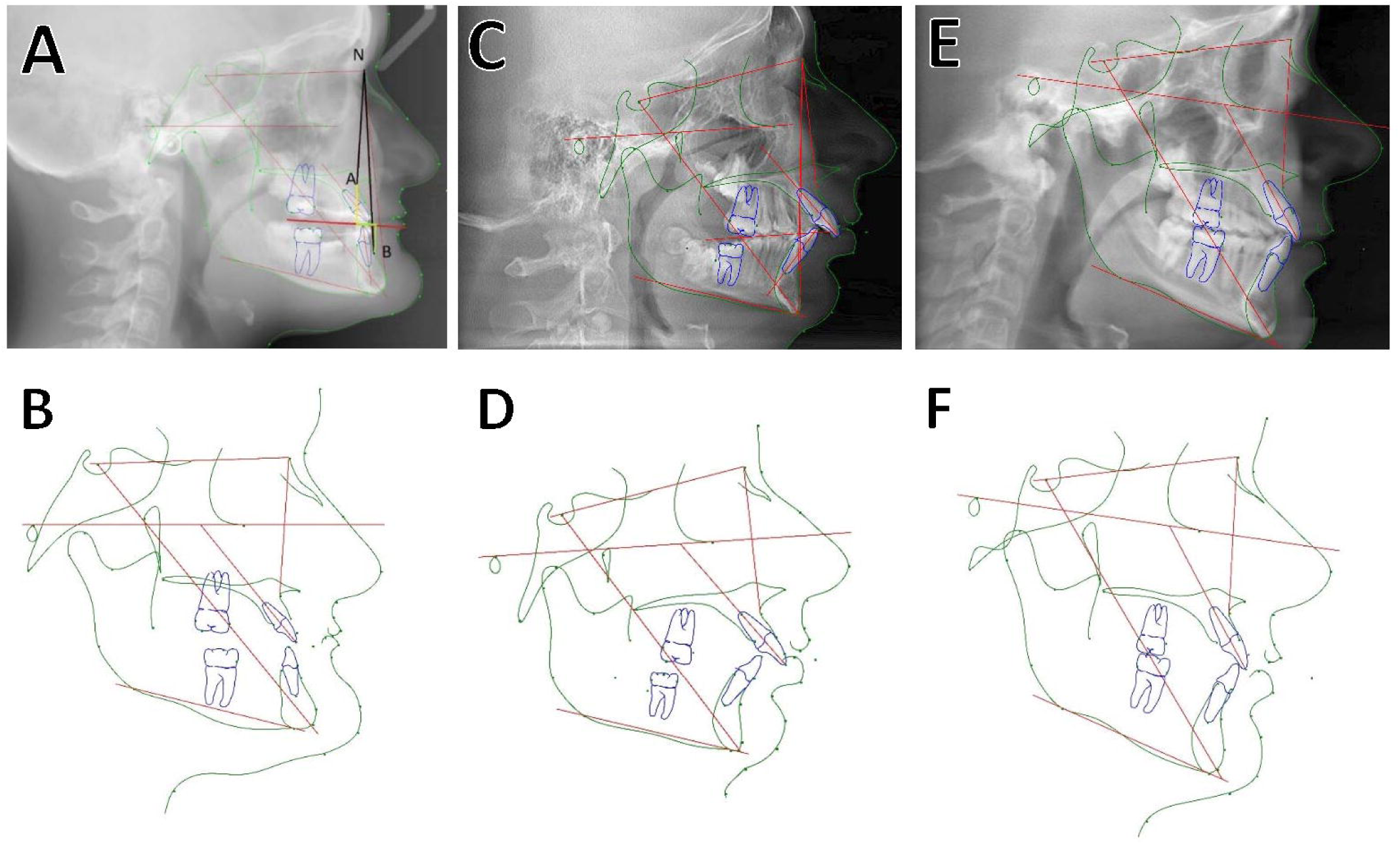
Cephalograms **A, B.** Individual 9 (girl, 14 years). The maxilla and mandible are significantly retrognathic. The ANB angle (-6.7 degrees, corresponding to 4.9 standard deviations [SD] below the reference mean value) and the Wits measurements (the length in mm of the yellow lines on the occlusal plane; 8.0 mm [3.1 SD above the reference mean value]) are abnormal, indicating a Class III biretrusive skeletal malocclusion compensated by severely proclined upper incisors (1 to NA line: 42.8 degrees; norm: 11 to 35 degrees) and severely retroclined lower incisors (lower incisor to mandibular plane angle: 67.8 degrees, norm: 83.4 to 99.4 degrees). The mandibular plane angle is also significantly decreased (15 degrees, norm: 25 to 41 degrees) indicating a severely brachycephalic facial type. **C, D.** Individual 10 (boy, 14 years). Cephalogram reconstructed from cone beam CT. The mandible is retrognathic (the SNB angle is 74 degrees, reference mean 78 [SD: 3.0] degrees). The maxilla-mandibular relationship is abnormal (the ANB angle is 7.4 degrees, reference mean 3.0 (SD: 2.0) degrees; Wits is 7.3, reference mean 2.0 [SD: 2.0] mm). All four second molars are impacted, the lower right first molar is semi-impacted and has only erupted partially. Roots on the lower left first molar are shortened. Typical Class II division 1 malocclusion with a retrognathic mandible and a reduced lower face height. **E, F.** Individual 13 (47 year old male). Almost normal cephalometric measurements.

CBCT scans were obtained from two study participants. An impacted upper second molar and the absence of the lower third molars were detected in Individual 11 (not shown). In Individual 10, the CBCT scan demonstrated impacted second molars and semi impacted lower first molars. An intervertebral disc calcification at the C2-C3 level was also identified (Figure 4).

**Figure 4.**
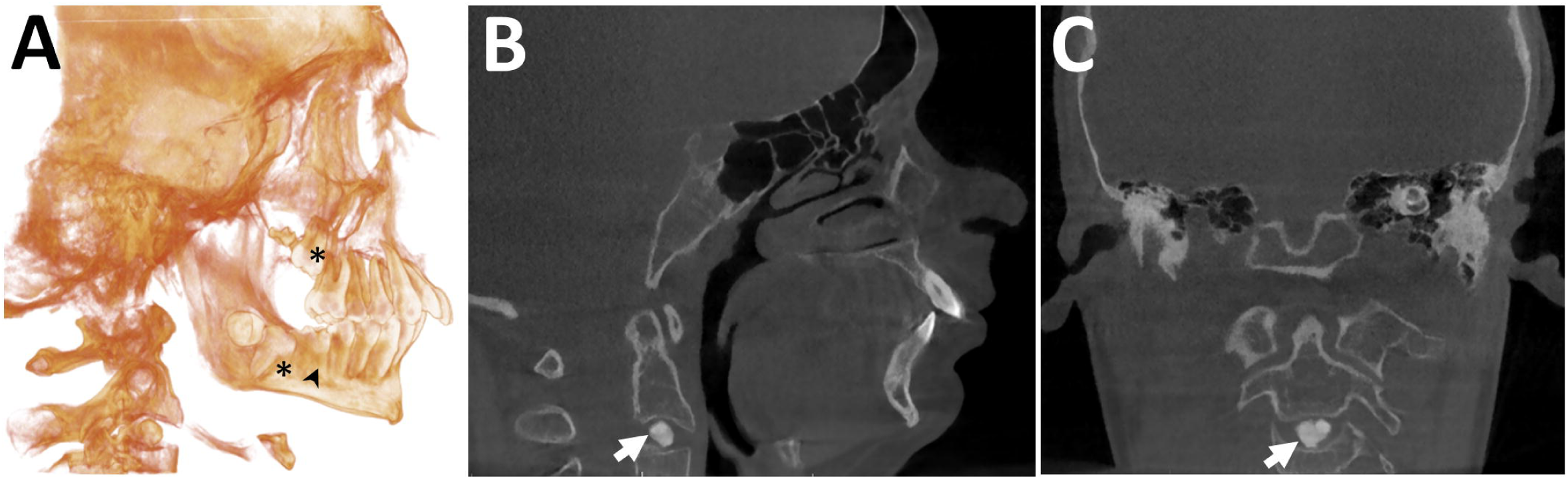
Cone Beam CT of Individual 10 (boy, 14 years). **A.** Three-dimensional volume rendering image. An unusual prognathic maxilla, impacted second molars (*) and lower right first molar is semi-impacted (arrowhead). **B.** Sagittal and **C.** Coronal images. There is a well-defined, round shaped radiopaque entity within the disc space at the C2-C3 level, consistent with intervertebral disc calcification (arrow). The adjacent vertebral end plates appear normal.

## Discussion

In this largest dental and craniofacial study population reported to date, we found that DI was absent but that missing teeth, especially premolars was a common occurrence. Malocclusion associated with either a retrusive maxilla or biretrusive jaws was also frequently observed. The facial type was either normal or brachycephalic and more than half of the individuals with OI type V presented with a retrusive (concave) profile and decreased lower face height due to underdeveloped maxillary and mandibular dentoalveolar processes.

The absence of DI in OI type V has been reported in the first description of the disorder (10) and has also been noted in subsequent case series (16, 17, 27). This could be explained by the lack of a collagen type I defect in OI type V, as collagen type I is the major protein of intertubular dentin (90%) (28).

Previous reports have noted that missing teeth are a common finding in OI caused by *COL1A1*/*COL1A2*, affecting the permanent dentition in 10% to 22% of individuals (4, 29). Premolars appear to be most commonly affected (4). The present study suggests that premolars are also most frequently missing in OI type V. Tooth development is a complex process involving signaling pathways that are also important for skeletal development such as WNT signaling (30). It is therefore intuitive that genetic defects leading to major abnormalities in bone cell function also affect tooth development, but the precise pathways whereby variants in *IFITM5* lead to tooth agenesis are unknown at present.

Previous studies have shown a high prevalence of class III malocclusion in the more severe OI types related to collagen type I mutations (OI types III and IV) (29, 31, 32). These malocclusions are caused by a severely hypoplastic maxilla and the counter rotation of the mandible resulting in severe negative overjet. Cephalometric studies suggest that the relative mandibular prognathism in these OI types in part reflects the decreased anterior length of the maxilla associated to a counterclockwise rotation of the mandible during growth (33). Our findings indicate that skeletal Class III malocclusion is less frequent in OI type V and that Class II malocclusion either of skeletal or dento alveolar origin is more frequent in OI type V than in other forms of moderate to severe OI. The severity of malocclusion varied widely in the present study independent of age. Six of the 14 study participants had severe malocclusion due to the presence of lateral open bite, multiple missing or impacted teeth and tooth migration. This indicates that their malocclusions presented significant therapeutic challenges where conventional orthodontic approaches would likely be inadequate. Nevertheless, the present study cohort presented with less severe malocclusion than what is typically seen in OI types III and IV caused by variants in *COL1A1* or *COL1A2* (5).

Regarding facial characteristics, reduced lower face height and concave profile were more common in our study cohort, suggesting that the dentoalveolar processes do not develop normally but do not result in the severely prognathic profile that is often observed in individuals with OI type II and IV. Deficient dentoalveolar development may be due to missing teeth altering the growth or may be a direct effect of the *IFITM5* variant.

In one of the two study participants who underwent CBCT scanning, we observed calcification of the intervertebral disk at the C2-C3 level. Calcification of intervertebral discs is a rare condition in children, which may present with neck pain, limited neck movement or may be discovered incidentally as in the present case. The cause of intervertebral disk calcification is unknown, but trauma and infection have been suggested as likely causes (34). Intervertebral disk calcification can lead to disc herniation, dysphagia or spinal cord compression (35), but can also resolve spontaneously (34). Thus, further follow-up of this individual will be important.

In conclusion, our study observed that OI type V is associated with missing teeth but not with DI. Facial analyses and cephalometric data show that OI type V is characterized by dentoalveolar alterations due to multiple missing teeth but not an exaggerated overjet, as is frequently observed in other types of OI. The malocclusion phenotype of OI type V varies widely, similar to the wide range of bone abnormalities found in the postcranial skeleton (16). The malocclusion of OI type V, if present, could be described as a biretrusive maxillomandibular malocclusion with reduced lower face height and multiple missing teeth. OI type V thus is associated with a unique pattern of craniofacial abnormalities that differ considerably from the malocclusion of other forms of moderate to severe OI.

## Acknowledgements

We are grateful to Jane Atkinson (National Institutes of Health) for helpful suggestions. This study was performed as an activity of the Brittle Bone Disease Consortium. The Brittle Bone Disease Consortium (1U54AR068069-0) is a National Center for Advancing Translational Sciences (NCATS) Rare Diseases Clinical Research Network (RDCRN), and is funded through a collaboration between the Office of Rare Diseases Research (ORDR), NCATS, the National Institute of Arthritis and Musculoskeletal and Skin Diseases (NIAMS), and the National Institute of Dental and Craniofacial Research (NIDCR). The content is solely the responsibility of the authors and does not necessarily represent the official views of the National Institutes of Health. The study was also supported by the Shriners of North America. None of the authors declares a conflict of interest.

